# UNSUPERVISED HARMONIZATION OF BRAIN MRI USING 3D CYCLE GANS AND ITS EFFECT ON BRAIN AGE PREDICTION

**DOI:** 10.1101/2022.11.15.516349

**Authors:** Dheeraj Komandur, Umang Gupta, Tamoghna Chattopadhyay, Nikhil J. Dhinagar, Sophia I. Thomopoulos, Jiu-Chiuan Chen, Dan Beavers, Greg ver Steeg, Paul M. Thompson, the Alzheimer’s Disease Neuroimaging Initiative (ADNI)

## Abstract

Deep learning methods trained on brain MRI data from one scanner or imaging protocol can fail catastrophically when tested on data from other sites or protocols - a problem known as *domain shift*. To address this, here we propose a *domain adaptation* method that trains a 3D CycleGAN (cycle-consistent generative adversarial network) to harmonize brain MRI data from 5 diverse sources (ADNI, WHIMS, OASIS, AIBL, and UK Biobank; total N=4,941 MRIs, age range: 46-96 years). The approach uses 2 generators and 2 discriminators to generate an image harmonized to a specific target dataset given an image from the source domain distribution and *vice versa*. We train the CycleGAN to jointly optimize an adversarial loss and cyclic consistency. We use a patch-based discriminator and impose identity loss to further regularize model training. To test the benefit of the harmonization, we show that brain age estimation - a common benchmarking task - is more accurate in GAN-harmonized versus raw data. *t*-SNE maps show the improved distributional overlap of the harmonized data in the latent space.

## 1. INTRODUCTION

Deep learning methods are now widely applied to brain MRI data for diagnostic classification, disease staging, and prognosis for a range of neurological and neurodevelopmental diseases. Even so, brain MRI protocols vary widely, and models trained on data from one scanner can fail when tested on data from a new site or protocol. To tackle this ‘domain shift’ problem, domain adaptation methods have been developed to adjust multisite brain MRI data to match a reference dataset or training data to facilitate machine learning on downstream tasks.

Two broad categories of domain adaptation methods have emerged: (1) adversarial methods that map source data into a site-invariant latent space [1, 2, 7], where features are optimized for the main task (e.g., disease detection) but also adapted to defeat an adversary that tries to predict which site the data came from; and (2) synthetic methods that also synthesize a new image to appear as if it came from another scanner, often using neural style transfer methods [4, 5]. Such reconstruction methods can also be extended to cross-modal data synthesis (e.g., simulating PET or CT scans from MRI) or for image enhancement with super-resolution [3].

Several GAN-based approaches have shown promising results for domain harmonization. For instance, Liu et al. [4] used style transfer methods to match brain MRI scans to a reference dataset. Sinha et al. [5] used attention-guided GANs for harmonization and demonstrated improvements in Alzheimer’s disease classification with harmonized data. Dinsdale et al. [2] developed a deep learning-based model to remove dataset bias while improving performance on a downstream task of brain age prediction.

Dewey et al. [6] presented DeepHarmony, a UNET-based architecture that requires a paired dataset for training. Many works limit the harmonization training to 2D slices rather than the whole 3D volume [2, 5, 6]. Zuo et al. [7] developed CALAMITI, which uses information bottleneck theory to learn a disentangled latent space that contains both anatomical and contrast information, enabling controllable harmonization in a parametrized protocol space.

Here we introduce an unsupervised CycleGAN method for domain adaptation which does not need any paired data across domains. We use 3D convolutions in our model’s generators and discriminators to harmonize full 3D MRI scans. We evaluate the harmonized scans by training machine learning models on a common benchmark downstream task (brain age estimation). We train downstream models for brain age estimation with harmonized and non-harmonized scans from five datasets. Our results illustrate improved brain age prediction after harmonization, suggesting that harmonization can improve the predictive performance of deep learning models in multisite predictive modeling.

## 2. DATASETS

**Table 1** summarizes the datasets analyzed in our study. We used 5 datasets: UK Biobank [8], ADNI [9], AIBL [10], OASIS-1 [11], and WHIMS [12]. Apart from WHIMS, we used 1,000 samples from each dataset.

**Table 1.**
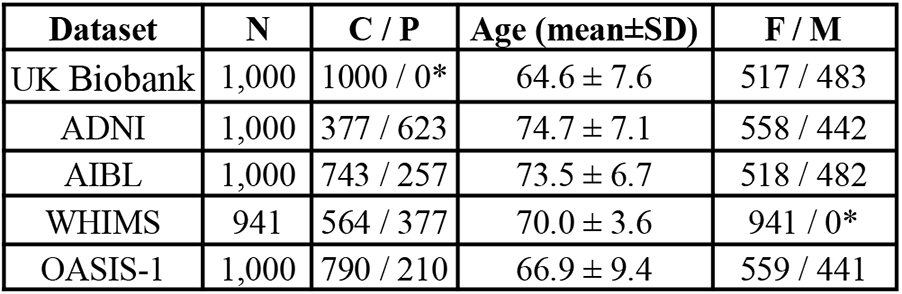
Statistics of the Training Dataset used for CycleGAN Training. N is the total number of samples. C, P, F, and M indicate the number of samples controls, patients, female, and male subjects, respectively. **\***WHIMS scans women only.

ADNI, AIBL, OASIS1, and WHIMS datasets include both controls and patient scans. The patients are diagnosed with Alzheimer’s disease in ADNI, AIBL, and OASIS1 datasets, and with Parkinson’s disease in the WHIMS dataset. UK Biobank has only controls. These datasets were used to train 4 CycleGAN models, mapping each of the four datasets (target datasets) ADNI, AIBL, OASIS, and WHIMS to (source dataset) UK Biobank and vice-versa.

All 3D T1-weighted MRI brain scans were pre-processed using a sequence of steps detailed in [13]. These included nonparametric intensity normalization (N4 bias field correction), skull stripping, 6 degrees-of-freedom registration to a template, and isotropic voxel resampling to 2 mm. Pre-processed images of size 91×109×91 were resized to 64×64×64, and intensities were linearly mapped to [0,1] using min-max normalization.

## 3. METHODS

Figure 1. summarizes our CycleGAN architecture. Similar to [14], it consists of two generators (G_X_: *X*→*Y* and G_Y_: *Y*→*X*) and two discriminators (D_X_ and D_Y_) for source domain *X* and target domain *Y*. With G_X_, we want to generate an image from the target distribution given an image from the source domain distribution and *vice versa* with G_Y_. We train the CycleGAN with an adversarial GAN loss and cyclic consistency loss. We use a patch-based discriminator and impose identity loss to regularize model training.

**Fig. 1.**
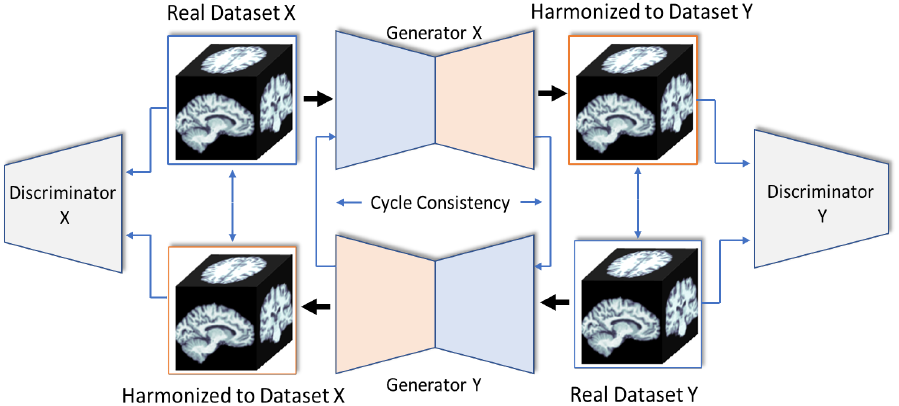
CycleGAN Architecture. Two generators and discriminators are trained to harmonize scans from dataset X to match the protocol of dataset Y and *vice versa*.

### 3.1 Model Architecture

The generator first transforms the input using a 3D convolution layer with 32 output channels and then passes it to the downsampling block. The downsampling block consists of two 3D convolution layers with 64 and 128 output channels that project the 64×64×64 input to 16×16×16. This is followed by 5 residual blocks, each having two 3D convolution layers with 128 output channels, and residual connections are made between each block [15]. Next, this output is passed to the upsampling block, which has two 3D transposed convolution layers with 64 and 32-channel outputs. Like UNET, we concatenate the downsampling block’s output with the upsampling blocks’ input. Finally, we concatenate the output of the upsampling block with the input image and pass it through a convolution layer to compute the output. We use instance normalization and ReLU non-linearity for all layers. For all convolution layers, we use padding and stride of 1, and kernel size 3 except for upsampling and downsampling blocks, for which stride was 2. The network’s output has the same dimensions as the input.

The discriminator uses a PatchGAN architecture [16, 17, 18] and has five 3D Convolution layers with 32, 64, 128, 256, and 1 output channel with kernel size 4. All layers except the first are followed by instance normalization and have a stride of 2, and all the layers use ReLU non-linearity except the last, which uses sigmoid activation. The output shape of the discriminator is 6×6×6 when the input dimension of the scan is 64×64×64.

#### 3.2.1 Adversarial Loss

We used the standard adversarial training to train the generator to synthesize real images. Instead of using the negative log-likelihood objective, we used least-squares loss, which provided stable training and better results. We use separate discriminators for each domain to compute the adversarial loss. The job of D_x_ is to distinguish between samples from the source data distribution, P(X), and the output of G_Y,_ and similarly, D_Y_ distinguishes between P(Y) and the output of G_X_. The overall adversarial loss is

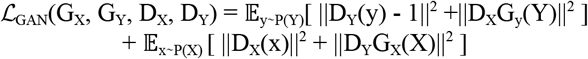

#### 3.2.2 Cycle-Consistency Loss

To reduce the space of possible mapping functions, we adopt the Cycle Consistency loss [14] which ensures that the two generators are cycle-consistent. That is, the scan translated to the target domain G_X_(x) and then translated back to the source domain - i.e., G_Y_(G_X_(x)) - must be mapped to the input from domain X. Effectively, we want G_Y_G_X_ to be identity mapping, i.e., x → G_X_(x) →G_Y_G_X_(x) ≈ x and similarly for G_x_G_Y_. We enforce this with pixel-wise L_1_ loss in both forward and backward directions, ensuring cycle consistency.

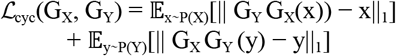

#### 3.2.3 Identity Loss

We further regularize the CycleGAN models by enforcing the identity loss similar to Taigman et al. [19], which showed that it helps to restrict the intensity range of the image for the image-to-image translation task. In the case of the grayscale scan, we noticed a slight improvement in the quality of the harmonized scans. The identity loss is computed as the pixel-wise L_1_ loss between the output of the generator (e.g., G_X_: *X*→*Y*) and the corresponding input (y∼P(Y)):

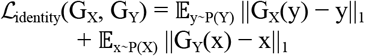

#### 3.2.4 Full Objective

We introduce additional hyperparameters, λ_1_ and λ_2_ to balance different loss terms. These are set to 10 and 0.1 during training. The overall objective loss for CycleGAN training is:

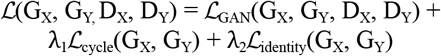

The patch-based adversarial loss ensures that the generator synthesizes images from distributions similar to the original distribution. The cyclic consistency and identity loss provide regularization on the generators.

### 3.3 Training

We trained 4 CycleGAN models, with *X* representing the UK Biobank source dataset and *Y* representing each of the ADNI, AIBL, OASIS, and WHIMS target datasets.

Each model was trained on pre-processed scans from the source and target datasets. Both generators and discriminators are trained with the Adam optimizer [20], with a learning rate of 10^−4^ and batch size of 4. The model was trained for 100 epochs with a multi-step learning rate scheduler with a gamma of 0.1 and steps on 35 and 75 epochs. Overall, our model has 16 million (M) parameters. Each generator and discriminator has 5M and 3M parameters, respectively.

## 4. RESULTS

We evaluated our domain adaptation model (1) via *t*-SNE for visual analysis of the data distributions, and (2) by testing if data harmonization reduced error on the downstream task of brain age estimation.

### 4.1 *t*-SNE and Clustering

Figure 2. visualizes the results of the harmonization of the source and target domains. Scans from the target domain were harmonized to the source domain. All the scans were resized to 8×8×8 before passing to the *t*-distributed Stochastic Neighbor Embedding algorithm to yield a 2D output for each scan. **Figure 2** shows the *t-*SNE embeddings of the unharmonized source and target domain. Without harmonization, source and target data are readily distinguished: in ADNI, all source domain points are clustered near the center, while target domain points are scattered around the source cluster. WHIMS data initially clustered almost entirely within the UK Biobank reference data. The second row shows *t-*SNE results after harmonizing the target to the source domain; there are now no distinguishable clusters, and source and target distributions overlap. This result is consistent with the goal that the source of the scans is hard to distinguish once they are harmonized.

**Fig. 2.**
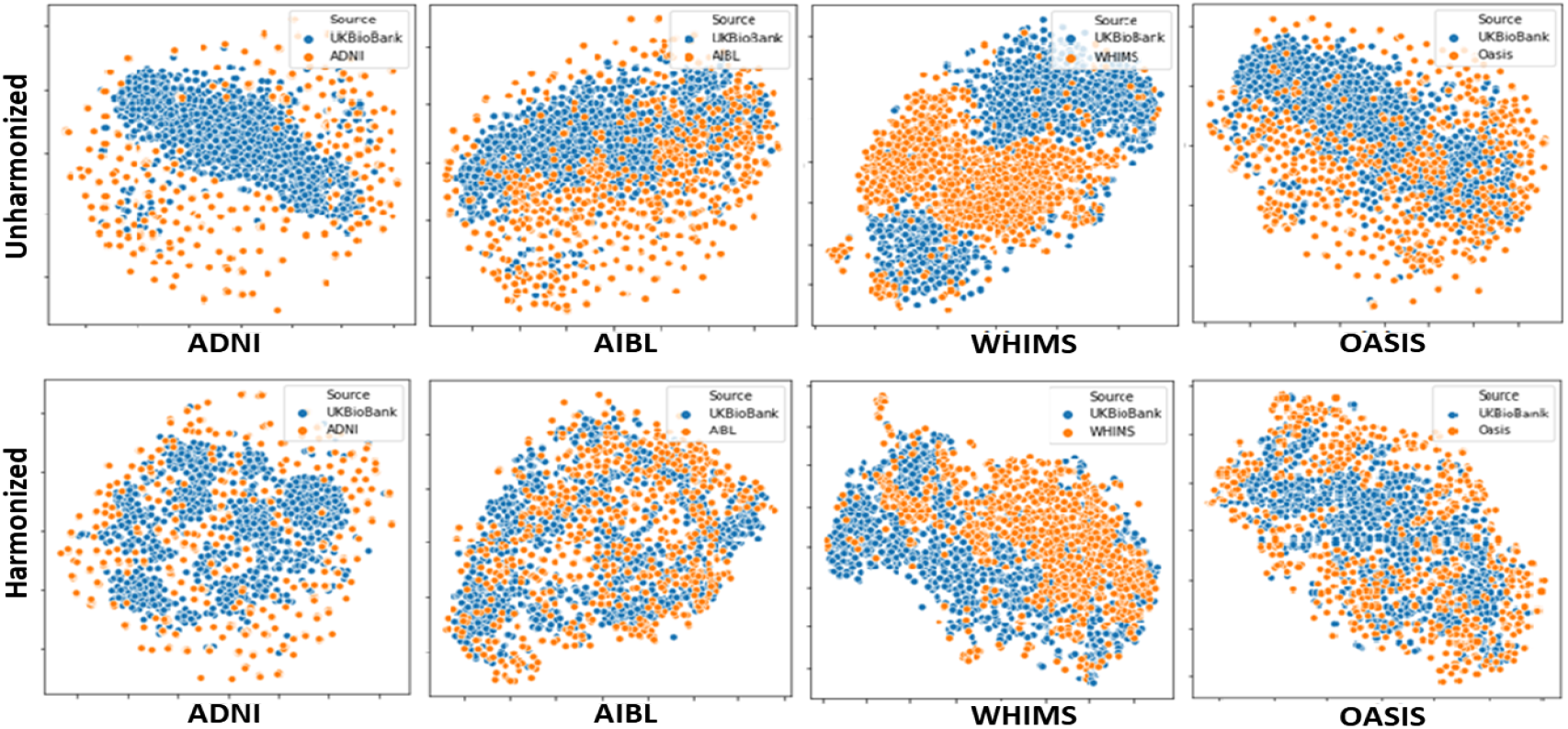
*t*-SNE visualization of harmonized (*bottom row*) and unharmonized (*top row*) scan from source and target domains. After CycleGAN harmonization, source and target scans are harder to distinguish in the *t*-SNE embedding

### 4.2 Brain Age Prediction

We tested how harmonization affects ‘Brain Age’ estimation, where a 3D CNN is trained on healthy controls’ MRI data to predict a person’s age from their scan. We used a model with 5 3D-Convolutional layers (output sizes: 32, 64, 128, 256, and 256), with a max pool in the first 3 layers. These layers were followed by LeakyReLU non-linearity and batch normalization. The last Conv. layer’s output was flattened and passed through 2 linear layers.

The CycleGAN can translate source datasets to target and target to source as well. Thus, when pooling the training datasets for the downstream task, we experimented with harmonizing all the target data to the source (UK Biobank in our case) and harmonizing UK Biobank data to the target datasets. We report the MAE score for the transformed dataset (a mixed hold-out test set from the source and the target) after applying harmonization in each direction. **Table 2** compares models trained on the pooled source and target datasets (a) with *no harmonization* (B, for *baseline performance*), or (b) after harmonizing the pooled data to either the source (To Source) or target (To Target).

**Table 2.** Brain Age Estimation on healthy controls used for training the unsupervised CycleGAN. S: subject’s ages, N: sample size. In the AIBL column, for example, we take 743 AIBL controls and 743 UKB controls, mix them, and split them into 1040 training and 446 test samples. After training, we report the MAE on the test samples with no harmonization (B, *baseline*). Then we harmonize all scans to ‘look like’ UKB and report the test MAE (to Source) and harmonize all scans to look ‘like AIBL’ (to Target) and report the test MAE.

|  | AIBL | ADNI | OASIS | WHIMS |
| --- | --- | --- | --- | --- |
| S | 72.9 ± 6.5 | 75.1 ± 6.5 | 65.37 ± 9.3 | 69.5 ± 3.6 |
| N | 743 | 377 | 790 | 564 |
| B | 4.557 | 3.437 | 5.109 | 3.856 |
| To Source | **4.250** | 3.772 | 5.487 | 3.645 |
| To Target | 4.551 | **3.149** | **4.770** | **3.552** |

In **Table 2**, N represents the total number of control subjects, which are split 70:30 ratio for training and evaluation. The datasets used in **Table 2** contain samples that were used to train the unsupervised CycleGAN models. This gives insights into using the same samples for unsupervised harmonization as well as the downstream task. We also report the results of training and evaluation on the samples which were not seen by CycleGAN in **Table 3** which gives us insights into the generalization capabilities of our harmonization approach.

**Table 3.** Brain age prediction results for ADNI and OASIS on a held-out set that were not used for training the CycleGAN. MAE is lower (better) after harmonization.

|  | ADNI |  | OASIS |  |
| --- | --- | --- | --- | --- |
| S | 76.4 ± 6.4 | | 65.4 ± 8.9 | |
| N | 1,000 |  | 742 |  |
|  | **Before** | **After** | **Before** | **After** |
| MAE | 4.387 | 3.929 | 4.344 | 4.339 |
| MSE | 30.539 | 25.428 | 29.780 | 31.231 |

**Table 2** shows improved brain age MAE on all 4 target domains after harmonization. We see similar trends with mean square error (MSE) but it is omitted due to space. For ADNI, WHIMS, and AIBL, we get better predictions when we translate from source to target when UK Biobank scans are harmonized to the target domain. For OASIS, harmonizing OASIS to UK Biobank was better. Visual insights after training at different numbers of epochs may be found on our GitHub repository (https://github.com/dheeraj-komandur/3D-CycleGAN-based-Harmonization).

**Table 3** reports brain age results for ADNI and OASIS datasets on a held-out set not used for training the CycleGAN. Prediction error changes little between **Tables 2 and 3**, suggesting that training on harmonized data can generalize to unseen samples.

## 5. CONCLUSION

We trained four CycleGAN models for deep learning-based harmonization of multicohort MRI, using the UK Biobank as the source domain and ADNI, AIBL, OASIS, and WHIMS as targets. Each CycleGAN model was trained with a joint adversarial loss, cycle consistency, and identity loss. We visualized the positive effects of harmonization in overlaying distributions by performing *t-*SNE on data before and after harmonization; clusters in the data distributions, found initially, were indistinguishable after harmonization. Further, brain age estimation improved in controls across all target domains after harmonization. In the future, we will assess if GAN-based harmonization improves multisite predictive modeling in AD and MRI-based amyloid level prediction. We also aim to train a model for multisite domain harmonization instead of 1-to-1 source-to-target mapping.

## 6. COMPLIANCE WITH ETHICAL STANDARDS

This study uses previously collected, anonymized, de-identified data. All the anonymized datasets are publicly available, including the UK Biobank (access approval #11559).

## 7. ACKNOWLEDGEMENTS

This work is supported by NIH (U01AG068057, P01AG055367).

